# Universal Single Copy Knock-In System in *Caenorhabditis elegans*: One Plasmid to Target All Chromosomes

**DOI:** 10.1101/2024.12.06.627295

**Authors:** Erica Dinneen, Carlos G. Silva-García

**Author notes:** Corresponding author (CGSG).

## Abstract

Successful transgenesis in model organisms has dramatically helped us understand gene function, regulation, genetic networks, and potential applications. Here, we introduce the universal single-copy knock-in system (Universal SKI System or U-SKI), designed for inserting any transgene by CRISPR/Cas9 in the *Caenorhabditis elegans* genome. The Universal SKI System takes advantage of a plasmid (pSKI), which can also be used for extrachromosomal arrays, to facilitate the insertion of a transgene at specific safe harbor loci on each autosomal chromosome. The pSKI plasmid contains multiple restriction sites for easy cloning and serves as a CRISPR/Cas9-based insertion repair template because it has two synthetic and long homology arms that recombine with the SKI cassettes. This system also uses a single crRNA guide, which acts as a Co-CRISPR enrichment marker. Overall, the Universal SKI System is highly flexible; with the same Universal SKI cassette on each autosome, researchers can select the insertion site and streamline tracking while reducing the complexity of expressing single-copy transgenes in *C. elegans*.

## INTRODUCTION

Diverse methods have been developed to express transgenes in *Caenorhabditis elegans*. Traditionally, exogenous genes have been introduced in muti-copy as extrachromosomal arrays^1^, gamma/UV integration^2^, or biolistic transformation.^3^ They can also be expressed in single-copy using MosSCI-based integration^4,5^. These techniques have certainly facilitated essential discoveries. However, the advent of CRISPR/Cas9 technology^6–8^ has revolutionized the way to modify the *C. elegans* genome, particularly for single-copy transgene expression.

CRISPR/Cas9 (clustered regularly interspaced short palindromic repeats-associated) enables the precise insertion of transgenes through homology-directed repair (HDR) that can label an endogenous gene at its native locus with a fluorescent protein^6,9–12^ or insert a transgene with its regulatory sequences.^13–16^ Currently, the insertion of exogenous genes remains a labor-intensive process. It involves several steps, including the preparation of the DNA template, the complexity involved in constructing the plasmids, and the use of co-selection or rescue markers linked to the transgenes. If these markers want to be removed, and if that is possible, the strain must be injected again or undergo multiple crossings. Moreover, there are limited genomic sites available for transgene insertion. Despite more than ten years of CRISPR/Cas9 research in *C. elegans*^8^, inserting exogenous genes and selecting appropriate protocols continue to be a challenge. To simplify the process of inserting single-copy transgenes into the *C. elegans* genome, we have developed the <u>U</u>niversal <u>S</u>ingle Copy <u>K</u>nock-<u>I</u>n System (Universal SKI System or U-SKI). Our innovative universal SKI system stands out for its remarkable simplicity: a single plasmid designed to target all chromosomes effectively. This streamlined approach not only enhances efficiency but also opens multiple possibilities for inserting exogenous DNA into the *C. elegans* genome.

Here, we present the universal SKI system, which consists of a collection of strains designed to insert a transgene using a single plasmid as a repair template. The universal SKI plasmid, pSKI, contains multiple restriction sites that facilitate the cloning of the gene of interest. This pSKI plasmid serves as a repair template for CRISPR/Cas9-based single-copy knock-in insertion at specific safe harbor loci on each autosomal chromosome. Since each autosome contains the same cassette, researchers have the flexibility to choose the final location for their gene of interest. Our universal SKI system smooths the process, offering a straightforward approach that minimizes complexity. It enables researchers to easily design their gene of interest, select chromosomal destinations, and track the genetic insertion efficiently, thereby reducing challenges associated with inserting single-copy transgenes.

## RESULTS

We sought to build a system that significantly simplifies the generation of single-copy transgenes in *C. elegans*. The universal SKI system consists of two parts. First is the plasmid, pSKI, in which any gene can be introduced by traditional cloning or Gibson reaction^17^ and used as a CRISPR/Cas9 repair template. Second, the universal SKI strains carry a cassette with identical sequences (homology arms) that flank the gene of interest in the pSKI plasmid. Each strain has one cassette per chromosome (I, II, III, IV, and V). The following sections describe how we built the universal SKI system, how it can be used, and its advantages and limitations.

### The plasmid

To engineer the universal pSKI plasmid that is easy to use even for less experienced researchers, we first designed a multi-cloning site sequence (MCS) (**Figure 1A**). This MCS hosts 47 single cuts, including the AscI and FseI restriction sites that have been amply used in *C. elegans* since these restriction sites do not exist in the nematode genome.^18,19^ In addition, the high number of single cuts in this MCS region will allow flexibility in designing and cloning any transgene or DNA sequence.

**Figure 1.**
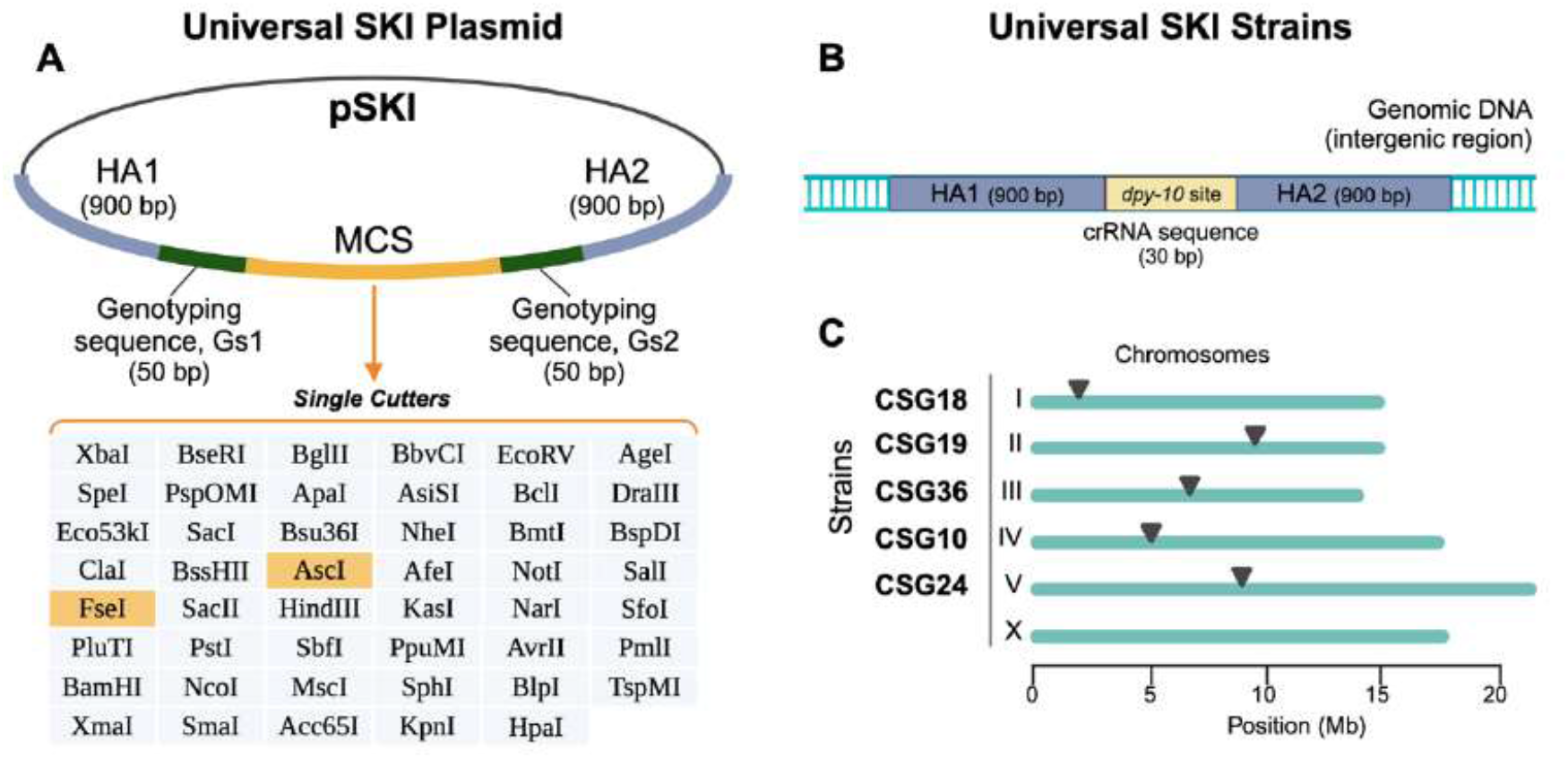
Generation of the Universal SKI System. (**A**) Schematic depiction of the pSKI plasmid. The pSKI is composed of two homology arms (HA1 and HA2, 900 bp each), two genotyping sequences (Gs1 and Gs2, 50 bp each), and a multi-cloning site sequence (MCS), which carries 47 single cuts. The table lists the restriction enzymes in 5’ to 3’ order. (**B**) Each Universal SKI strain carries the *dpy-10* site for the *dpy-10* crRNA, which is flanked by the same HAs that are present in the pSKI plasmid. (**C**) Every SKI cassette was introduced into a defined autosomal location by CRISPR/Cas9. Details about the construction of the Universal SKI System can be accessed in the Methods section.

To use the pSKI plasmid as a CRISPR template and enhance the efficiency of homologous recombination^7,20^, we specifically designed two synthetic homology arms (HAs). Each HA comprises 900 bp (∼50% GC content), which does not exist in the *C. elegans* genome. Furthermore, the HAs do not overlap with any *C. elegans* genomic region to prevent off-target silencing by potential endogenous RNA interference, which occurs when a few identical segments of at least 25 nucleotides exist within the gene.^21^ We designed the HAs for single-copy insertion by homologous recombination, and the gene of interest (GOI) will reside between them: HA1::GOI::HA2. Since the GOI significantly differs for each single-copy insertion and experiment needs, we added two small sequences for universal genotyping during the PCR screening (genotyping sequences: Gs1 and Gs2). Each Gs has 50 bp, does not exist in the *C. elegans* genome, and flank the MCS with the final design as HA1::Gs1::MCS::Gs2::HA2 (**Figure 1A**). In summary, the researcher can use the MCS region to open the pSKI plasmid and introduce the gene of interest using their preferred cloning method, obtaining the following design: HA1::Gs1::**GOI**::Gs2::HA2.

### The strains

Having the universal pSKI plasmid, we sought to generate transgenic *C. elegans* strains in which a single copy of the HAs has been knocked in at a defined safe harbor locus. These strains also contain the target sequence of a well-characterized crRNA that could later be used to knock in the GOI by CRISPR/Cas9 and, simultaneously, be used as a Co-CRISPR marker (**Figure 1B**).

To define safe harbor loci for the universal SKI system, we used those that we have previously characterized as safe harbor loci. We developed a single-copy knock-in loci for defined gene expression (SKI LODGE) system that allows rapid single-copy tissue-specific expression of any gene under five different promoters: ubiquitous (*eft-3p* -*eef-1A*.*1p*-), neuronal (*rab-3p*), muscle (*myo-3p*), germline (*pie-1p*), and intestinal (*elt-2p*) promoters.^14^ We showed that the insertion of the SKI LODGE cassettes in these regions gives stable expression with no silencing.^14^ The strains also have no defects in fertility, embryonic lethality, developmental timing, and lifespan.^14^ Taking advantage of these well-characterized safe harbor loci, we inserted the universal SKI cassettes in these regions using the cloning-free and *in vitro* assembly CRISPR/Cas9 approach.^14,22,23^

We generated a suit of transgenic strains with an equal cassette design. Each universal SKI strain consists of the two exact 900-bp HAs present in the pSKI plasmid (**Figure 1A and B**), and between them is a highly efficient CRISPR target sequence copied from the *dpy-10* gene: HA1::*dpy-10* site::HA2 (**Figure 1B**). By inserting a 30-base protospacer and PAM sequence from *dpy-10* gene^24^, hereafter referred to as “*dpy-10* site”, we can simultaneously induce double-stranded breaks at both 1) the universal SKI cassette *dpy-10* site and 2) the endogenous *dpy-10* locus using a single crRNA guide. Other studies, including our own, have demonstrated that this approach is highly efficient and cost-effective, requiring only one crRNA guide.^14,25^ In order to introduce the *dpy-10* site into the universal SKI cassette, we used another well-established and identifiable Co-CRISPR target gene, *dpy-5*.^14^ The combination of long HAs with the high efficiency of the *dpy-10* site amplifies the likelihood of inserting a template into the Universal SKI cassette. We have generated five universal SKI strains, with the universal SKI cassette introduced into defined intergenic regions in all autosomal chromosomes (I, II, III, IV, and V) (**Figure 1C**). We engineered all universal SKI strains following the CRISPR/Cas9 protocol based on *in vitro* assembly and using PCR products as templates.^14,22,23^ We did one CRISPR edit step to introduce the final cassette using two overlapping PCR fragments (see Methods). Finally, all universal SKI end lines were outcrossed six times to eliminate potential off-targets. Since the same cassette is harbored on all autosomal chromosomes, the user can select the chromosomal destination based on their GOI and cross different universal SKI lines into each other.

### Testing the Universal SKI System

To verify that our universal SKI system could be used to introduce an entire gene in one step to drive single-copy expression, we 1) cloned a fluorescent reporter gene into the universal pSKI plasmid and 2) inserted the reporter gene by CRISPR knock-in, using the pSKI as a repair template.

We selected a strong fluorescent reporter to test our universal SKI system. Using the Gibson assembly protocol^17^, we cloned the following transgene into the pSKI plasmid: *myo-3p::mCherry::unc-54 3’ UTR* (see Methods) (**Figure 2A**). Without including the HAs, the total length of this fluorescent reporter is 4091 bp and drives mCherry expression in wall muscles. CRISPR/Cas9 mix was assembled *in vitro*^14^ using this pSKI-reporter plasmid as a repair template (**Figure 2B**). As expected, utilizing only one crRNA guide, we obtained *dpy-10* mutants (Co-CRISPR marker) that also showed muscular expression of mCherry in all our universal SKI strains (**Table S1**). Because not all GOI carry a fluorescent reporter to track or the fluorescence is not strong enough to be detected under a standard dissecting UV microscope, we specifically designed our universal pSKI plasmid with two sequences (Genotyping sequence 1 and 2, Gs1 and GS2) for screening purposes: HA1::**Gs1**::GOI::**Gs2**::HA2 (**Figure 1A**). These Gs1 and Gs2 sequences are always introduced along with the GOI and serve as genotyping sequences for both 5’ and 3’ ends (**Figure 2C**). Each Gs1 and Gs2 region amplifies a fragment around 1000 bp, which is used to screen dumpy animals for single-copy insertion by PCR (**Figure 2D, screening**). We have designed and tested primers that can be used to screen the universal SKI strains (see the step-by-step guide in **File S1**). We have also designed and tested primers to outcross the final universal SKI lines that carry the GOI into the N2 wild type (**Figure 2D, outcrossing**). Since all our universal SKI strains harbor the same cassette, having these “universal” primers simplifies the screening and process of generating single-copy transgenes.

**Figure 2.**
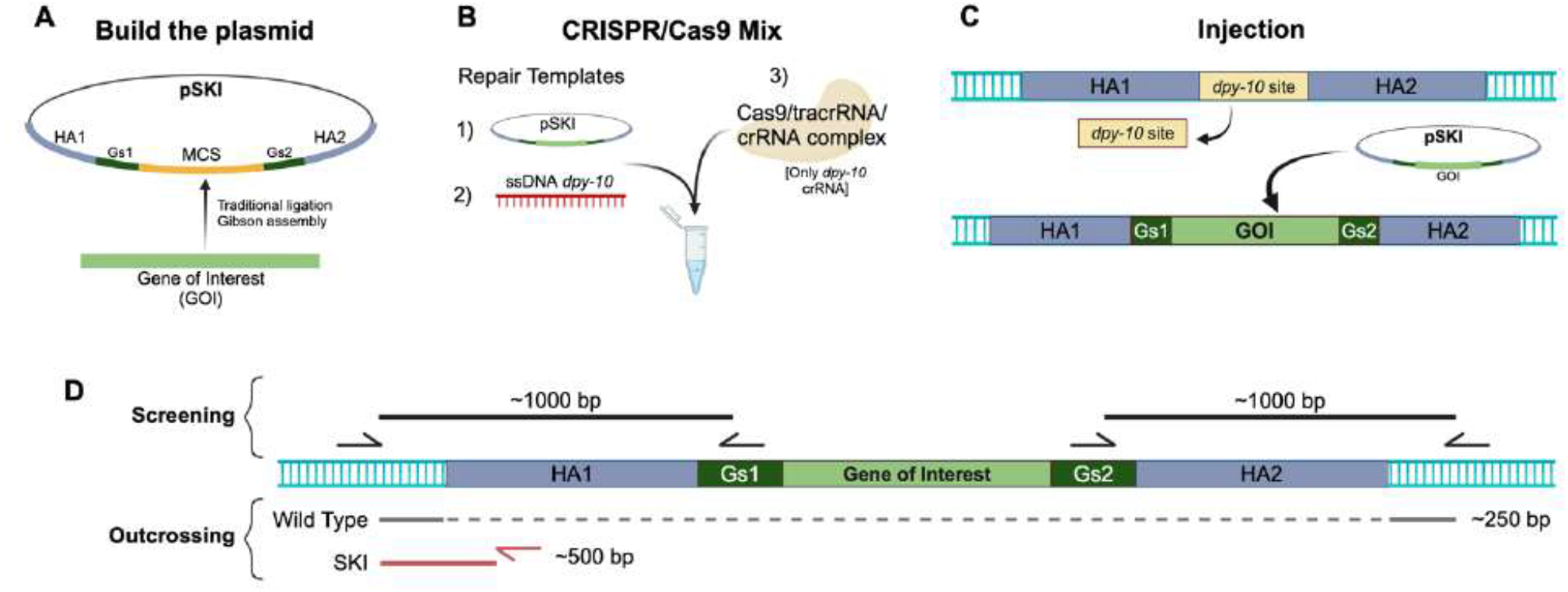
Workflow for using the Universal SKI System to generate single-copy insertions. (**A**) The gene of interest (GOI) is cloned into the pSKI plasmid. (**B**) Assemble CRISPR/Cas9 mix *in vitro* using the pSKI plasmid as a repair template. (**C**) Injection of this mix replaces the *dpy-10* site with the GOI. (**D**) Using the universal SKI primers, the insertion of the GOI is screened either in the 5’ region (genotyping sequence 1, Gs1) or the 3’ region (genotyping sequence 2, Gs2). After identifying and confirming the GOI integration, another set of primers is used to outcross the SKI strain. For details about the cloning, insertion, and screening, see the step-by-step user guide in **File S1**.

We used the PCR strategy for the Gs1 and Gs2 regions to discern whether the *myo-3p::mCherry::unc-54 3’ UTR* transgene is expressed as extrachromosomal arrays or as a single copy (**Figure 2D, screening**). We successfully analyzed the 5’ and 3’ ends by PCR, with a final single-copy insertion percentage of 3.0%, 8.8%, 3.8%, 4.7%, and 3.9% for chromosomes I, II, III, IV, and V, respectively (**Table S1**). Overall, our universal SKI system is simple and allows easy single-copy insertions. The universal pSKI plasmid allows the researcher the flexibility to generate any desired transgene, and, as an added value, it can be easily shared and edited among the *C. elegans* community. The universal SKI strains offer the option to choose the target chromosome to insert the GOI. Finally, the efficiency of CRISPR editing depends on several factors (some of them discussed here^10,26^). In the case of our universal SKI system, we have found that achieving a successful SKI insertion depends on the purity of the CRISPR mix, a well-honed microinjection technique, and a clean PCR screening.

## DISCUSSION

Here, we present the universal SKI toolkit for generating single-copy insertions into the *C. elegans* genome at specific safe harbor locations. We engineered the pSKI plasmid to function as a repair template for CRISPR/Cas9-based insertions. Additionally, we developed a set of universal SKI strains that allow the pSKI plasmid to be inserted. Thus, the performance of our system relies on its simplicity: it uses just one plasmid for all constructions that can be inserted in any autosomal chromosome.

What does our universal SKI system contribute to the existing CRISPR-based options for expressing single-copy transgenes in *C. elegans*? Several techniques have been developed to express single-copy transgenes in the nematode genome. Among the most recent transgenic methods include the use of Flp, Cre, and phiC31 recombinases^27–29^, Split-wrmScarlet and split-sfGFP^12^, modular safe-harbor transgene insertion (MosTI)^16^, and the Mos1-mediated Single-Copy Insertion (MosSCI) method, which is widely used to establish stable transgenic strains.^4,5,30,31^ All these methods have been beautifully designed and have certainly advanced the generation of single-copy *C. elegans* transgenic lines. However, *C. elegans* researchers usually must decide whether to use efficient methods to integrate single-copy transgenes at random sites or less efficient and laborious methods to target a specific genomic site. Thus, our toolkit complements the current transgenic methods and provides some important advantages. We have specifically developed the universal pSKI plasmid to be simple. It can be used for even less experienced researchers due to the plasticity of the multi-cloning site sequence (MCS) (**Figure 1A**). Further, our universal SKI strains are manageable to inject as they behave as wild type and are precisely designed not to cause disruptions in the *C. elegans* genome since we synthetically designed the homology arms (HAs), and they are integrated into specific safe harbor loci. Having the universal SKI cassette in all autosomal chromosomes gives freedom to select the desired targeted region (**Figure 1B**). Lastly, the universal SKI primers that target the genotyping sequences (Gs1 and Gs2) streamline the PCR screening, although the researcher needs to corroborate the entire insertion at the end. Overall, we aimed to develop a simple and straightforward transgenic method for *C. elegans* researchers.

Finally, the most crucial advantage of our universal SKI system is its flexibility. Since our pSKI plasmid backbone is built on the widely used *C. elegans* plasmid for extrachromosomal arrays, the pPD95.75 plasmid^32^, the pSKI plasmid can also be used for extrachromosomal arrays. In another scenario, suppose a researcher has their gene of interest (GOI) already cloned into a plasmid for extrachromosomal expression. In that case, a simple digestion and ligation can extract the GOI and introduce it into the pSKI plasmid. **Table 1** lists the advantages and potential combinations that can be applied along with our system. All universal SKI reagents are available freely to the *C. elegans* community, and a step-by-step user guide is included in **File S1**. Strains reported here, new strains, updated protocols, and all sequences can be found at https://www.theSGlab.com/resources.

**Table 1:**
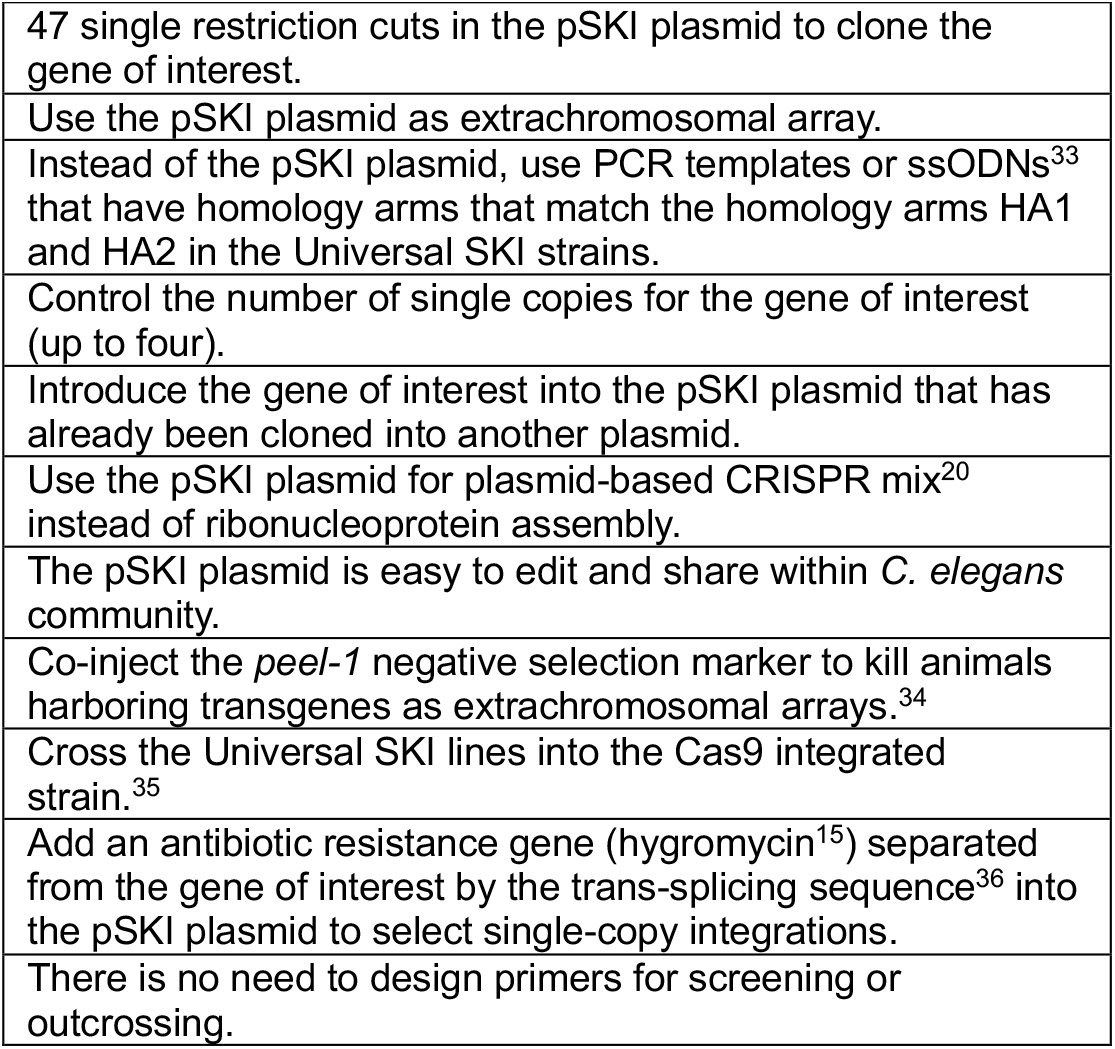
Advantages and potential combinations for the Universal SKI System.

### Limitations of the study

While the Universal SKI System offers advantages over other CRISPR-based options for expressing single-copy genes, some experimental limitations remain. There is no SKI cassette in chromosome X, which could be a strain of interest to study X-chromosome dosage compensation.^37^ The SKI cassette is in all autosomal chromosomes, but more strains are needed in order to provide at least two options for each chromosome. Big insertions are always a challenge. Although our system expedites this process, we are currently generating new strains that can contribute to inserting long exogenous sequences by carrying more than one site for double-strand breaks. This approach has been shown to enhance the insertion of large DNA fragments. However, it deletes hundreds of genomic DNA^10^. Our future universal SKI strains will solve this issue: no deletion of genomic DNA, use only one crRNA guide to induce 2-3 double-strand breaks, and, importantly, these strains will use the same pSKI plasmid described here. This means genes already cloned into the universal pSKI plasmid can be integrated into the new strains as a single copy.

## MATERIAL AND METHODS

### Strains and maintenance

Bristol N2 strain was provided by the Caenorhabditis Genetics Center (CGC), which is funded by NIH Office of Research Infrastructure Programs (P40 OD010440). The worms in this experiment were grown on nematode growth medium (NGM) plates. These plates were seeded with OP50-1, an *Escherichia coli* bacterial strain. The *C. elegans* used in this study were maintained under standard procedures.^38^

### Strains and sequences generated in this work

The strains generated in this study are listed below in **Table 2** and will be available for distribution through the CGC. All sequence files for the universal SKI lines are available at https://www.theSGlab.com/resources. All crRNAs, tracrRNA, and oligonucleotides are listed in **Tables 3 and 4**.

**Table 2.**
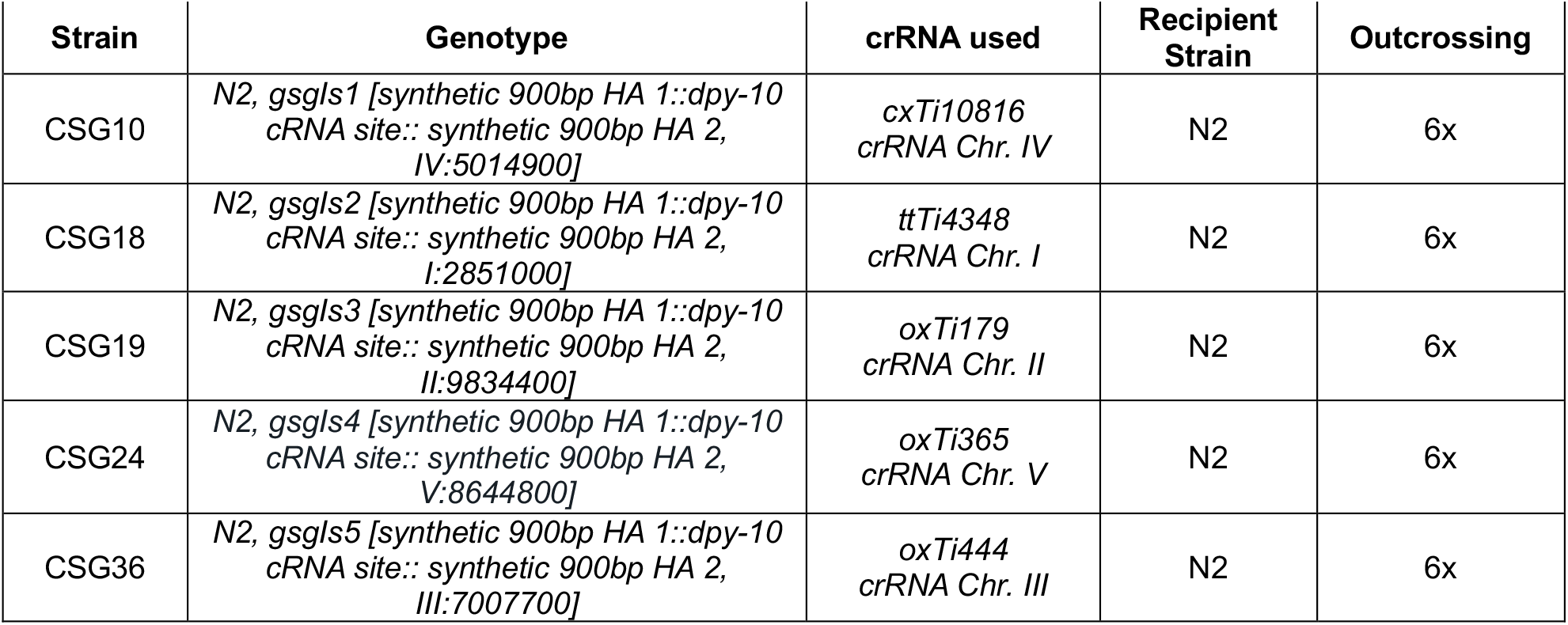
Strains generated in this study.

**Table 3.**
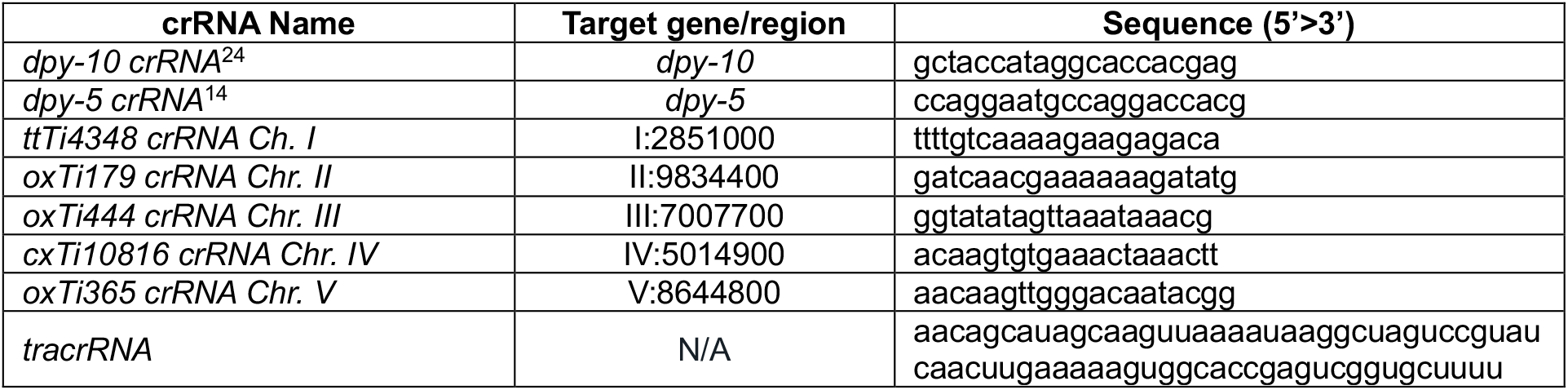
crRNAs and tracrRNA used in this study.

**Table 4.**
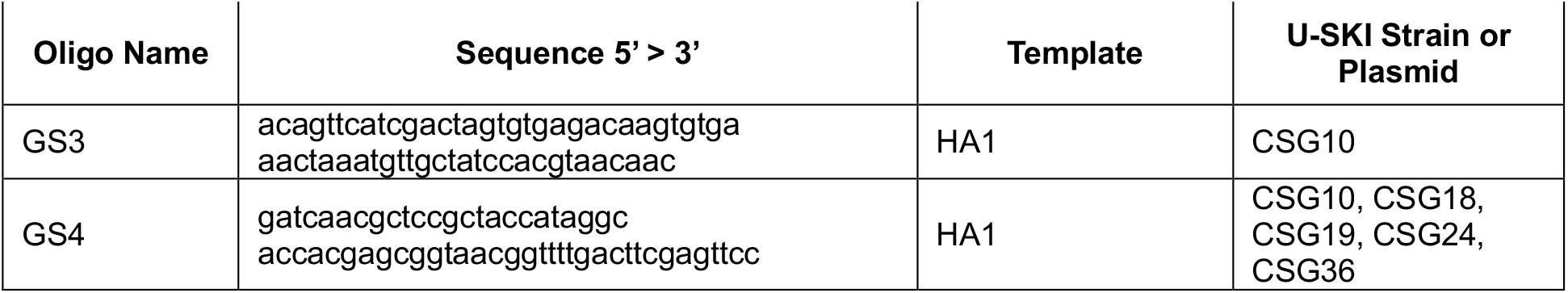
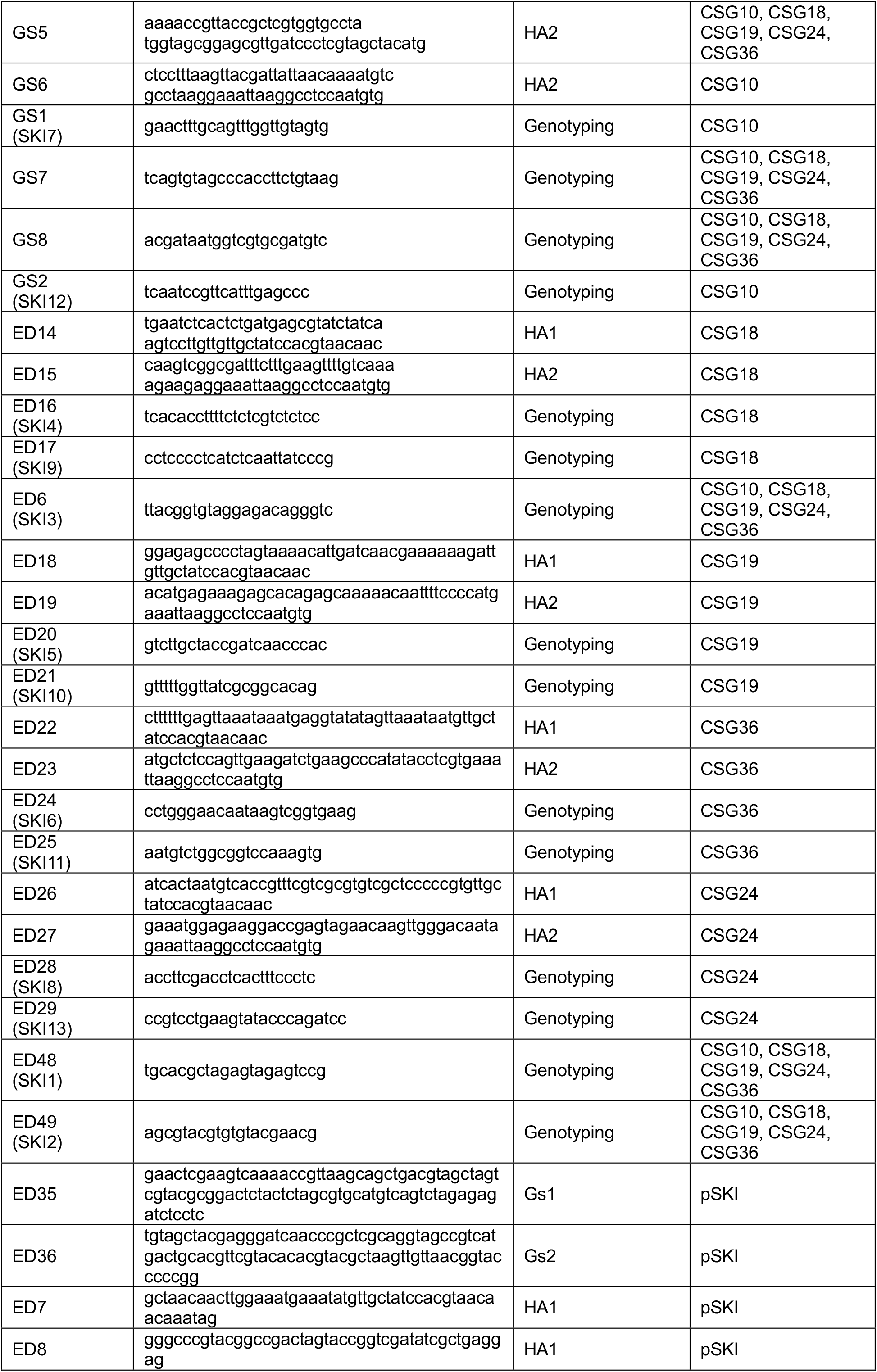
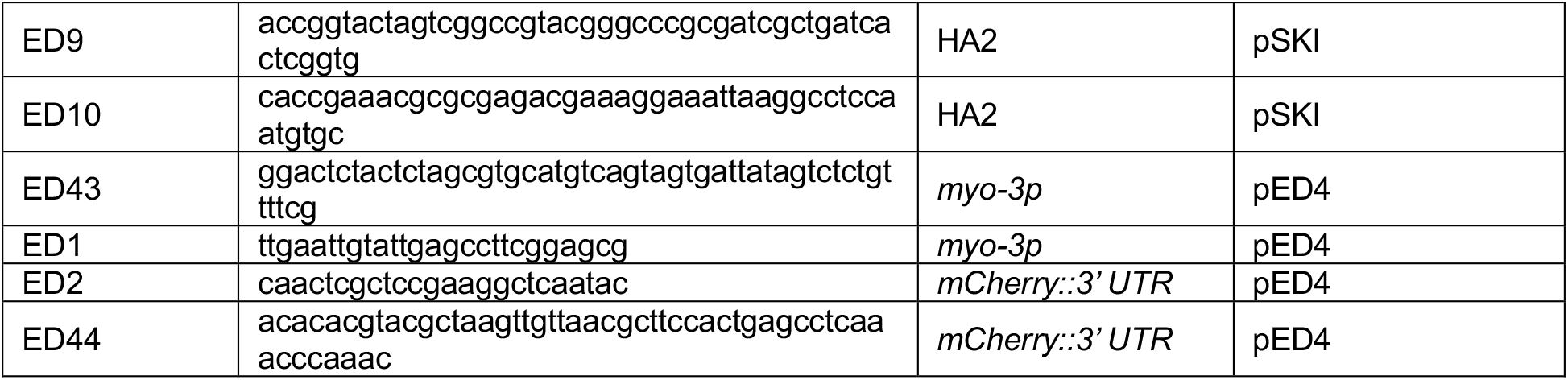
Oligonucleotides used in this study.

### Identification of harbor loci

We utilized previously characterized safe harbor loci to knock in the universal SKI cassette. These sites have been well characterized and tested in our previous SKI LODGE system.^14^ The universal SKI system cassettes were inserted at the following positions: Chr. I:2851000, Chr. II:9834400, Chr. III:7007700, Chr. IV:5014900, and Chr. V:8644800.

### Generation of the pSG (pSKI) plasmid

The universal SKI plasmid consists of two synthetic homology arms (HAs), two genotyping sequences (Gs1 and Gs2), and a multi-cloning sequencing site (MCS). First, we designed the two synthetic HAs. Each HA comprises 900 bp with around 50% GC content and was generated using the random DNA sequence generator.^39^ After each sequence was generated, they were run for Basic Local Alignment Search Tool (BLAST) in order to guarantee the sequences were not found anywhere else in the *C. elegans* genome, thereby preventing unintended off-target silencing. A similar approach was followed to generate the synthetic Gs1 and Gs2 sequences. The MCS, which resides between Gs1 and Gs2, contains 47 single cuts. The sequences for the HAs and MCS were ordered from Genewiz as single-stranded DNA templates. Gs1 and Gs2 templates were generated via PCR utilizing primers listed in **Table 4**. The plasmid pPD95_75 served as the backbone for the pSKI plasmid. pPD95_75 was digested with HindIII and ApaI (NEB). The digested plasmid was purified using a Zymo Gel DNA Recovery kit (D4007). The single-stranded DNA templates and backbone were ligated following standard Gibson Assembly protocol^17^ and transformed into NEB 5-alpha competent *E. coli* (NEB C2987H). The final pSKI plasmid was sequenced to corroborate the editions. pPD95_75 was a gift from Andrew Fire (Addgene plasmid #1494).

### Cas9 purification

Cas9-His-tagged was purified by overexpressing in *E. coli* BL21 Rosetta cells. Nickel-NTA beads were used to pull it down followed by HPLC purification. Briefly, we inoculated LB culture with SpCas9 containing *E. coli* Rosetta cells and agitated for 3-6 h until the culture reached an OD of 0.6-0.8. The culture was cooled down on ice water to below 20 ºC, and 0.5 mM IPTG was added, followed by incubation overnight in a shaker at 20 ºC. The culture was spun, and the bacterial pellet was resuspended in 27.5 ml of 50 mM Tris pH 8, 150 mM NaCl, 10% glycerol and 2 mM TCEP. Cells were lysed by adding 2 ml 10X FastBreak buffer (Promega V8571) and 5 μl Benzonase, incubated for 15-20 min at RT, spun at 38000g for 15 min at 4 ºC, and the supernatant was collected. Ni-NTA resin was washed in an equilibration buffer (50 mM Tris pH 8, 500 mM NaCl and 10% glycerol). The supernatant was then incubated with 1 ml of this pre-washed Ni-NTA resin at 4 ºC for 1 h. The sample was poured into a disposable column and washed with wash buffer (20 ml of 50 mM Tris pH 8, 500 mM NaCl and 10% glycerol, 20 mM Imidazole and 2 mM TCEP). The sample was then eluted using elution buffer containing 5 ml 50 mM Tris pH 8, 500 mM NaCl and 10% glycerol, 400 mM Imidazole, and 2 mM TCEP. The eluted Cas9 protein was diluted 1-2 fold with 1X PBS and loaded onto a 5 ml heparin column equilibrated with 1X PBS. The sample was eluted in a linear salt gradient from 0.1 to 1 M NaCl in 1X PBS and 1 ml fractions were collected. Fractions were analyzed on a gel and pooled all Cas9 containing fractions followed by concentrating them down to 3 ml volume with Amicon Ultra-15 filter (30 kDa cutoff, spun 4000 g 20 min at 4 ºC). We exchanged the buffer for 1X PBS, 10% glycerol, and 2 mM TCEP using a PD-10 column. The insoluble protein was spun down at 4 ºC for 15 min and concentrated down to 4/12, a final volume of 500 μl. Purified Cas9 was filter sterilized (0.22 μm), and the final concentration was measured by Nanodrop and stored at -80 ºC.

### CRISPR/Cas9 mix and microinjection

All CRISPR edits and insertions required to generate and test the universal SKI strains were performed using the CRISPR protocol adapted from Paix *et al*. 2015^22^ and Silva-García *et al*. 2019.^14^ Briefly, homology repair templates were amplified by PCR, using primers that introduced a minimum stretch of 35 bp homology to the universal SKI cassette at both ends. CRISPR injection mix reagents were added in the following order: 0.375 μl Hepes pH 7.4 (200mM), 0.25 μl KCl (1M), 1.25 μl tracrRNA (8 μg/μl), 0.5 μl dpy-10 crRNA (8 μg/μl) or 0.5 μl dpy-5 crRNA (8 μg/μl), 0.5 μl dpy-10 ssODN (1000 ng/μl) or 0.5 μl dpy-5 ssODN (1000 ng/μl), and PCR repair template(s) up to 400 ng/μl final in the mix. Water was added to reach a final volume of 8 μl. 2 μl purified Cas9 (12 μg/μl) was added at the end, mixed by pipetting, spun for 2 min at 13000 rpm, and incubated at 37 ºC for 10 min. Mixes were microinjected into the germline of day 1 adult hermaphrodite worms using standard methods.^2^

### Universal SKI cassette construction

All universal SKI strains were generated by the abovementioned CRISPR protocol, using at least 35 bp of homology as recombination arms. Two templates were used to generate the universal SKI lines. The templates were amplified from the pSKI plasmid. One template consisted of HA1 and the other of HA2. The genomic *dpy-10* site was added between the two templates: HA1::*dpy-10* site::HA2. Each template, including homology arms for the corresponding genomic region and the overlapping sequence for the *dpy-10* site, with around 980 bp in length. The primers used to amplify the HA1 and HA2 and the specific crRNAs for each strain can be found in **Table 3 and 4**. In order to introduce the *dpy-10* site into the universal SKI strains, we used another easily identifiable Co-CRISPR target gene, *dpy-5*^14^. All final universal SKI strains were verified by sequencing and were outcrossed to N2 at least six times to remove the Co-CRISPR marker mutation (*dpy-5*) as well as any additional unwanted off-site mutations.

### Verification of Universal SKI strains

To validate the universal SKI strains, we generated a transgene reporter based on our universal SKI system. The following transgene was cloned into the pSKI plasmid: *myo-3p::mCherry:unc-54 3’ UTR*. The transgene was amplified from the pCFJ104 plasmid (Addgene plasmid #19328) by PCR. Briefly, the pSKI plasmid was digested with XbaI and KpnI (NEB). Then, two PCRs were performed, one for amplifying the *myo-3p* and one for *mCherry::3’ UTR*. The primers utilized in these PCRs are listed in **Table 4**. These two purified PCRs and the digested pSKI plasmid were ligated following standard Gibson Assembly protocol and transformed into NEB 5-alpha competent *E. coli* (NEB). After verification, the final plasmid, pED4, was injected into each universal SKI strain, following the CRISPR protocol from Silva-García *et al*. 2019^14^ (see above CRISPR/Cas9 mix section).

Immediately after injection, individual worms were placed at 20 ºC. 3-4 days after injection, all plates were screened for *dpy* animals and red fluorescent worms. Worms expressing mCherry were then lysed and underwent PCR for both homology arms (HA1 and HA2) to determine single-copy insertions. F1 worms were placed in 5 µl of single worm lysis buffer (30 mM Tris pH 8.0, 8 mM EDTA pH 8, 100 mM NaCl, 0.7% NP-40, 0.7% Tween-20, and 100 µg/ml proteinase K) and lysed for 1 h at 60 °C, followed by incubation at 95 °C to inactivate the proteinase K. We then screened for the CRISPR edit by PCR using Apex Taq RED Master Mix 2.0X (Genesee Scientific) as recommended by the manufacturer, using 1 µl of worm lysate as a template. Animals were genotyped utilizing the genotyping SKI primers. A more detailed approach regarding genotyping can be found in our step-by-step guide in File S1.

## Ethics statement

*C. elegans* is not protected under most animal research legislation

## ACKNOWLEDGMENTS

We thank all Silva-García laboratory members for their helpful discussions. The graphics in Figures 1 and 2 were created with biorender.com. CGSG is funded by NIA/NIH R00AG065508 and the American Federation for Aging Research.

## AUTHOR CONTRIBUTIONS

CGSG designed the study. ED generated the final pSKI plasmid. ED generated and tested all universal SKI alleles used in this study. ED and CGSG wrote the manuscript.

## SUPPLEMENTAL MATERIAL

**Table S1.**
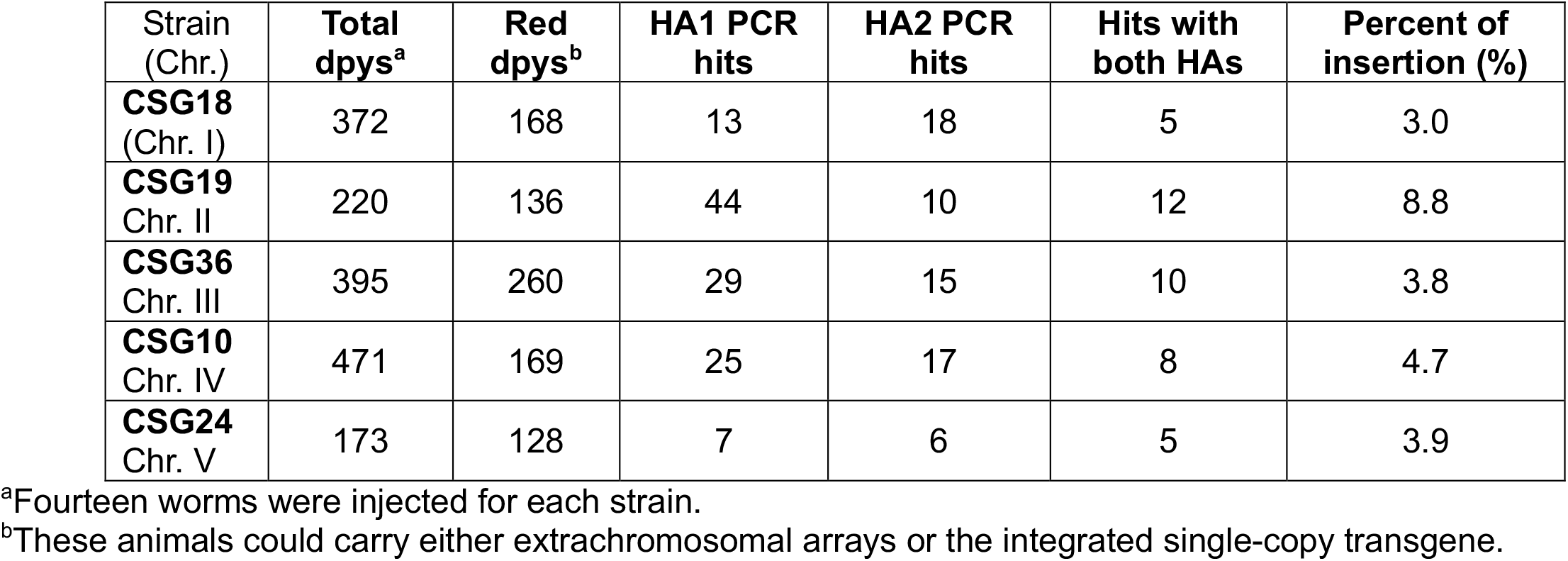
Homology-directed repair (HDR) efficiencies in the Universal SKI strains.

### File S1

Step-by-step guide for using the Universal SKI System

Strains reported here, new strains, updated protocols, and all sequences can be found at https://www.theSGlab.com/resources

### Step 1

Clone your transgene of interest (GOI) in the pSKI plasmid by traditional cloning or Gibson reaction.

➢ HA: Homology arms (1 and 2); 900 bp each.
➢ Gs: Genotyping sequence (1 and 2); 50 bp each.
➢ MCS: Multi-cloning site sequence (47 cuts); 222 bp.
➢ AscI and FseI restriction sites (not found in the *C. elegans* genome).
➢ pSKI: 4728 bp and Amp resistant.

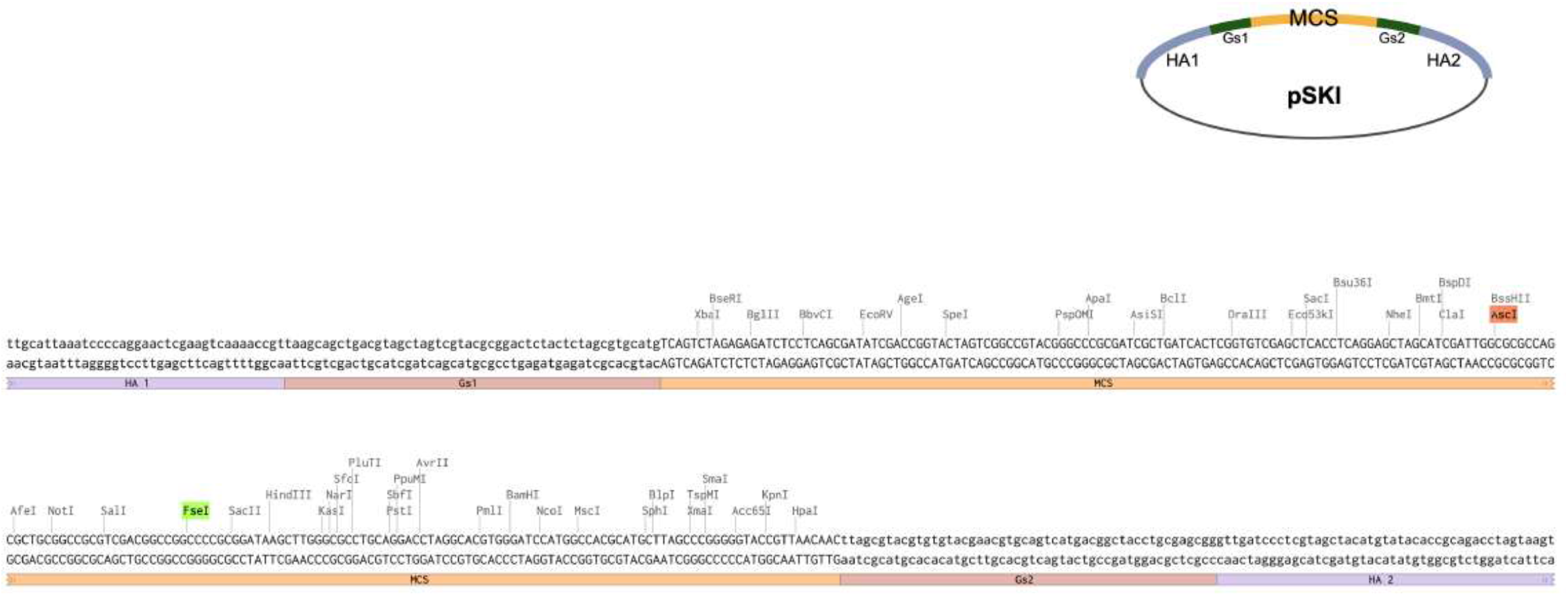

### Step 2

Select the Universal SKI strain to knock in your gene of interest.

➢ All strains have the following cassette:

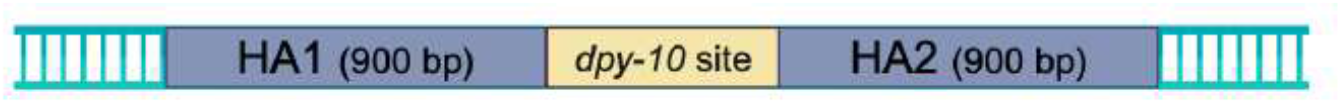

➢ Strains and chromosome positions:

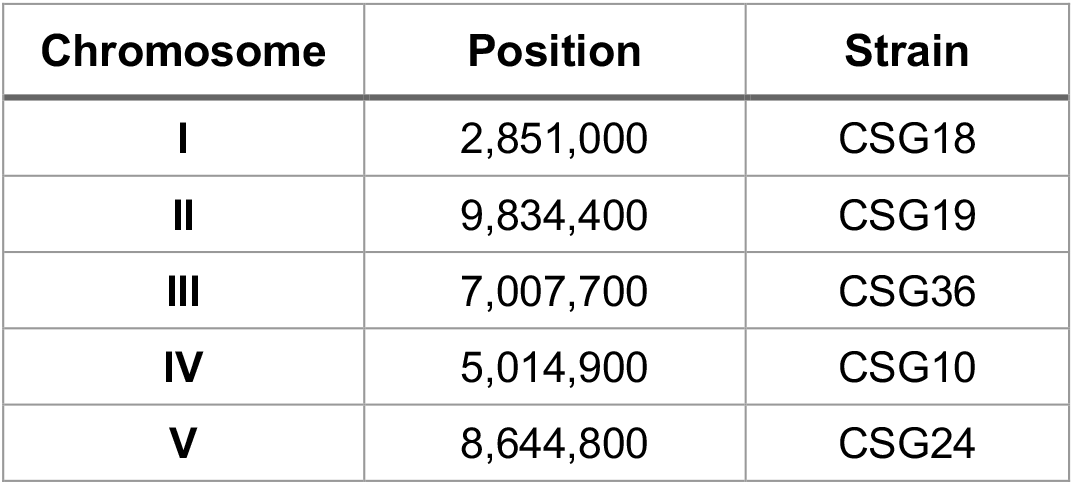

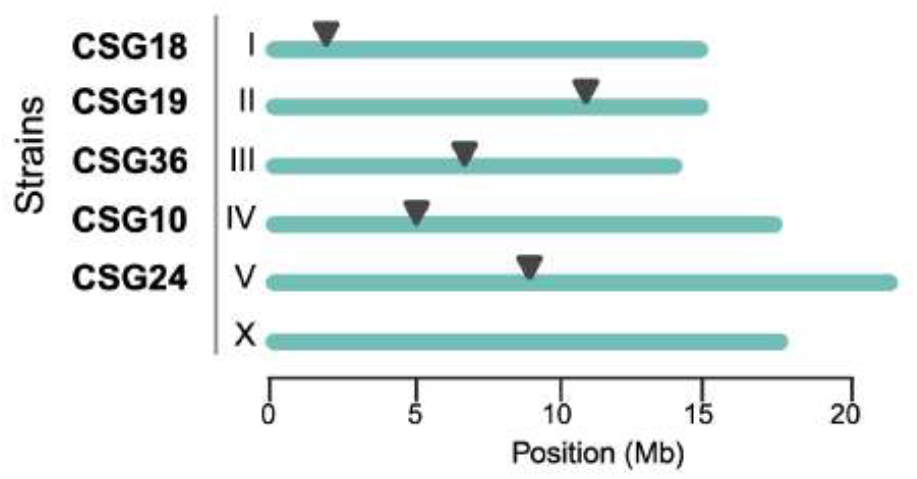

### Step 3

Assemble of CRISPR/Cas9 complex *in vitro*:

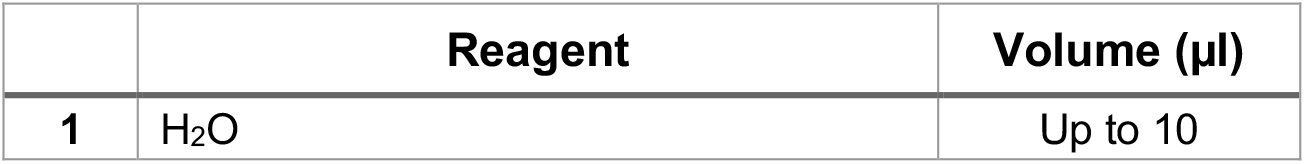
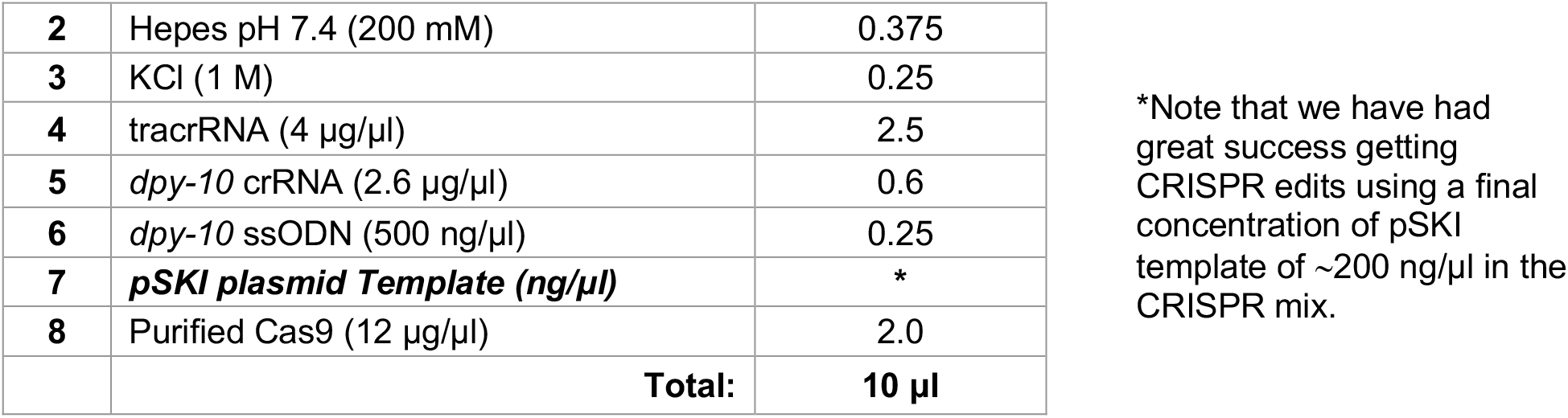

➢ Injection mixes can be prepared in advance without Cas9 protein, separated into 2 tubes of 4 μl, and stored at -80ºC.
➢ We have found that frozen mixes stored for up to one year at -80ºC are still effective at generating CRISPR edits.
➢ Before injecting, thaw one 4 μl mix, add 1 μl of purified Cas9 (12 μg/μl), mix by pipetting, spin for 2 min at 13000 rpm, and incubate at 37ºC for 10 min.
➢ Inject into day 1 adults of the relevant Universal SKI strain.

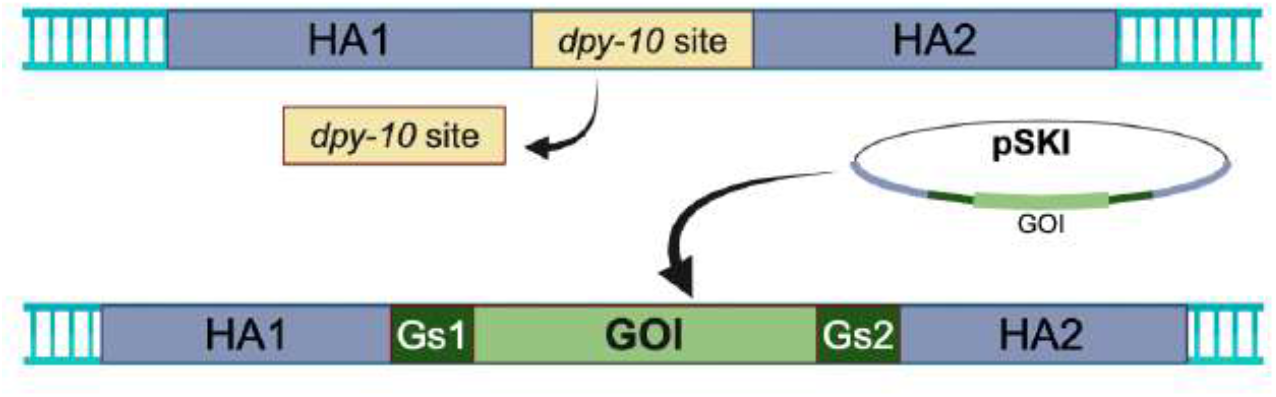

### Step 4

3-4 days after injection, screen for plates that have produced many dumpy and/or roller progenies.

➢ From these plates, individually plate single *dpy* animals and allow them to lay eggs before screening them for your desired edit. Screening can be done by looking for fluorescent protein expression and/or by genotyping them by PCR.
➢ We have designed two specific sequences (Gs1 and Gs2) for PCR genotyping.
➢ We have also annotated and tested additional Universal SKI Primers to facilitate the screening process.
➢ The screening consists of two independent PCRs:

1. PCR 1: Amplify the 5’ region using the Gs1 sequence.
2. PCR 2: Amplify the 3’ region using the Gs2 sequence.

➢ **Note**: If you design new primers for genotyping, avoid designing them inside the HA1 or HA2 regions since these homology arms sequences are also present in the pSKI plasmid. Then, design primers inside of your gene of interest.
➢ Use the following model, table, and gels as a reference for PCR screening.

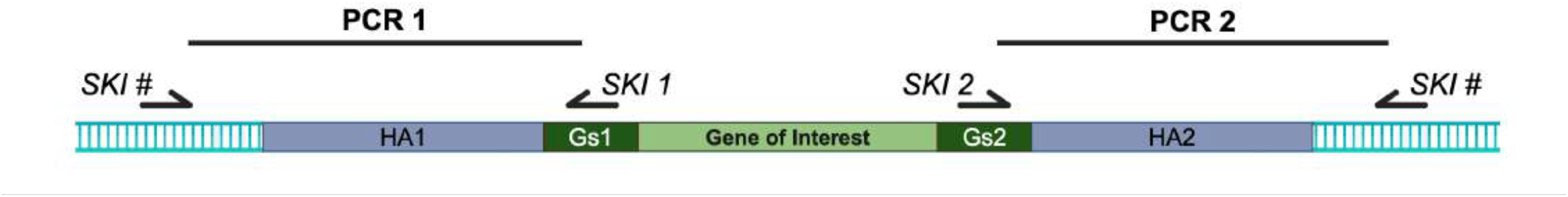

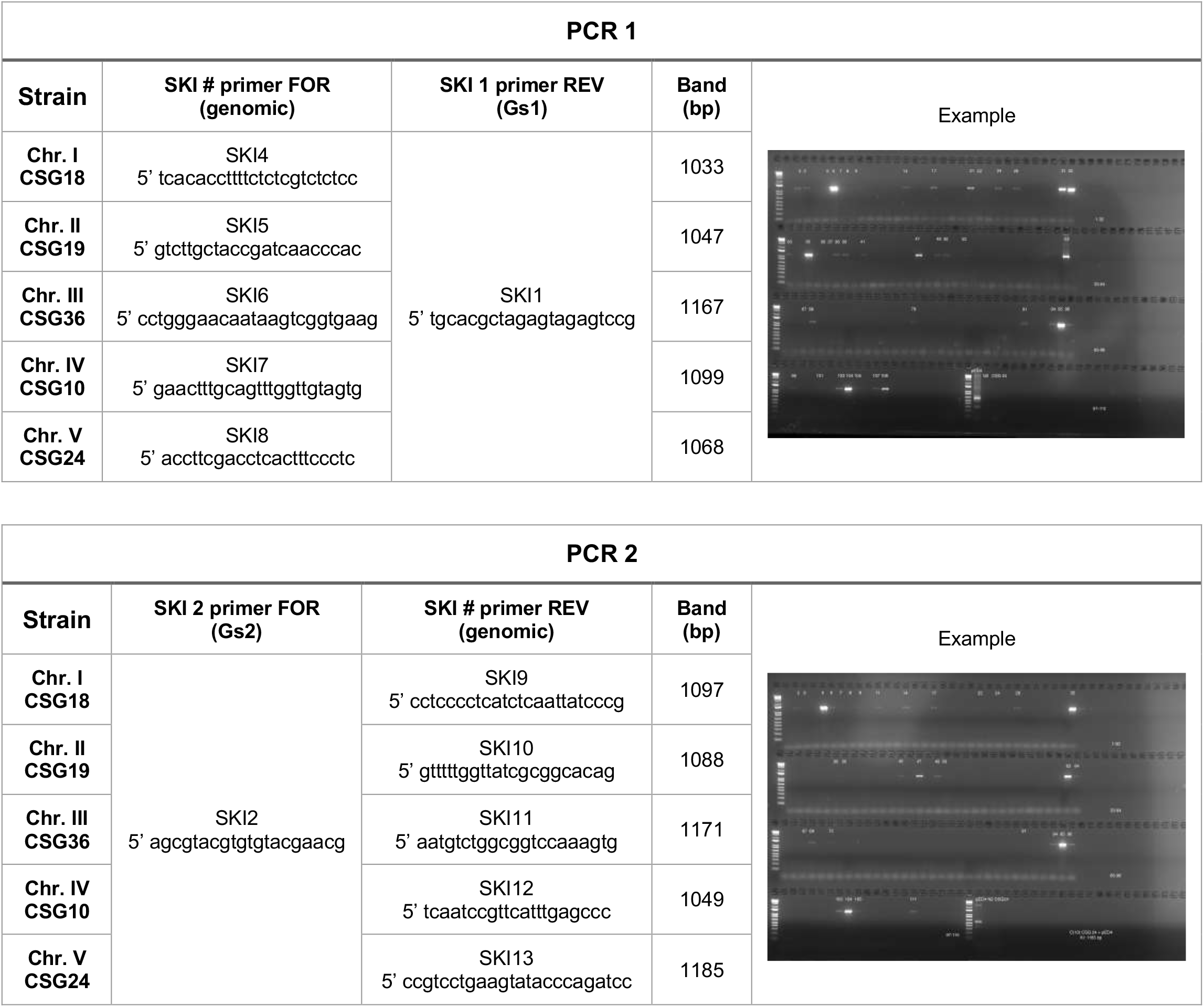

➢ PCR conditions:

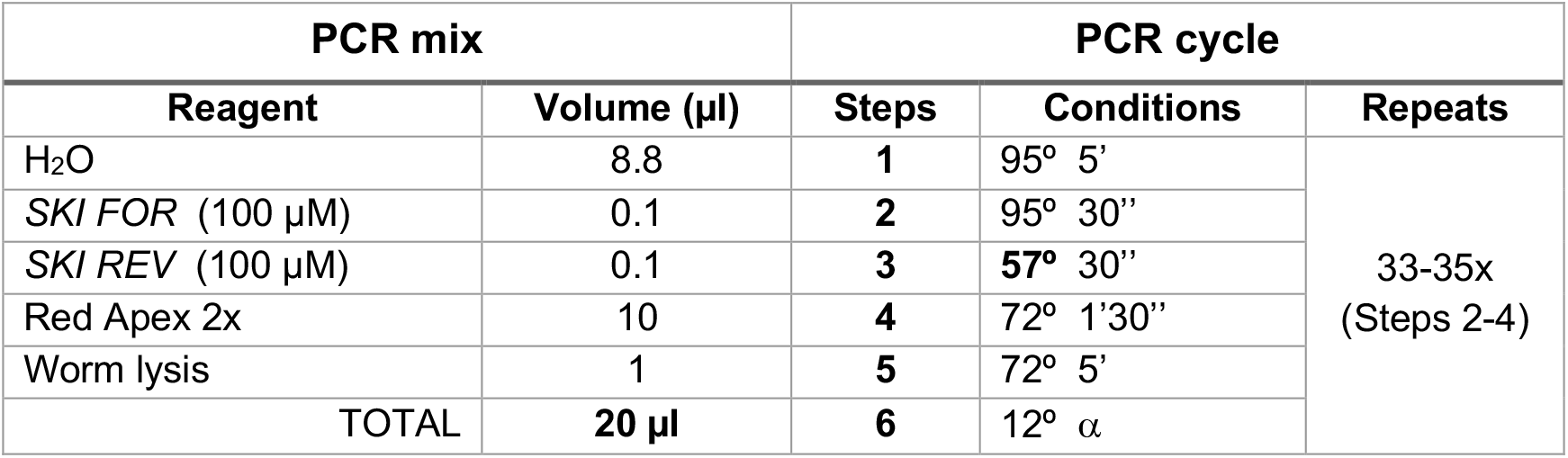

### Step 5

From those F1 animals that screen positive for **both** PCR1 (5’ region) and PCR2 (3’ region), go back to the plates they were picked from and pick 4-6 of their F2 progeny to individual plates, allow them to lay eggs, and then screen them as above. Continue doing this until getting homozygous animals for your desired edit. Confirm that the knock-in CRISPR edit is correct by sequencing.

### Step 6

Outcross your newly made Universal SKI line to eliminate the *dpy-10* Co-CRISPR edit.

➢ Note that the outcrossing will use the wild-type N2 strain, not the Universal SKI strains.
➢ The following model and table show primers flanking the Universal SKI cassette that can be used to outcross the new SKI line to N2.
➢ Note that the external primers are the same as those used for the CRISPR screening.
➢ Follow the same PCR conditions described above and reduce the extension to 1 minute.

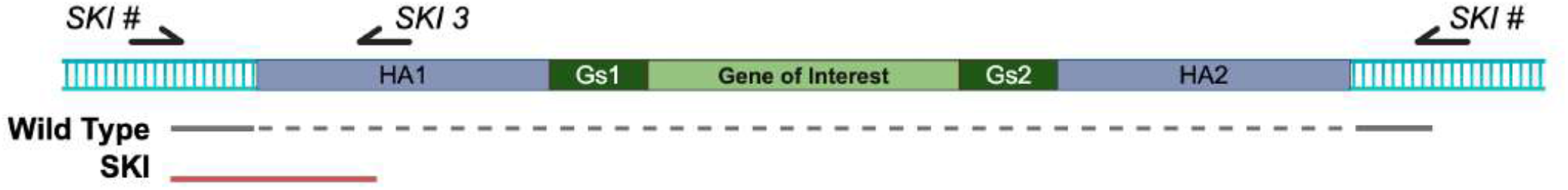

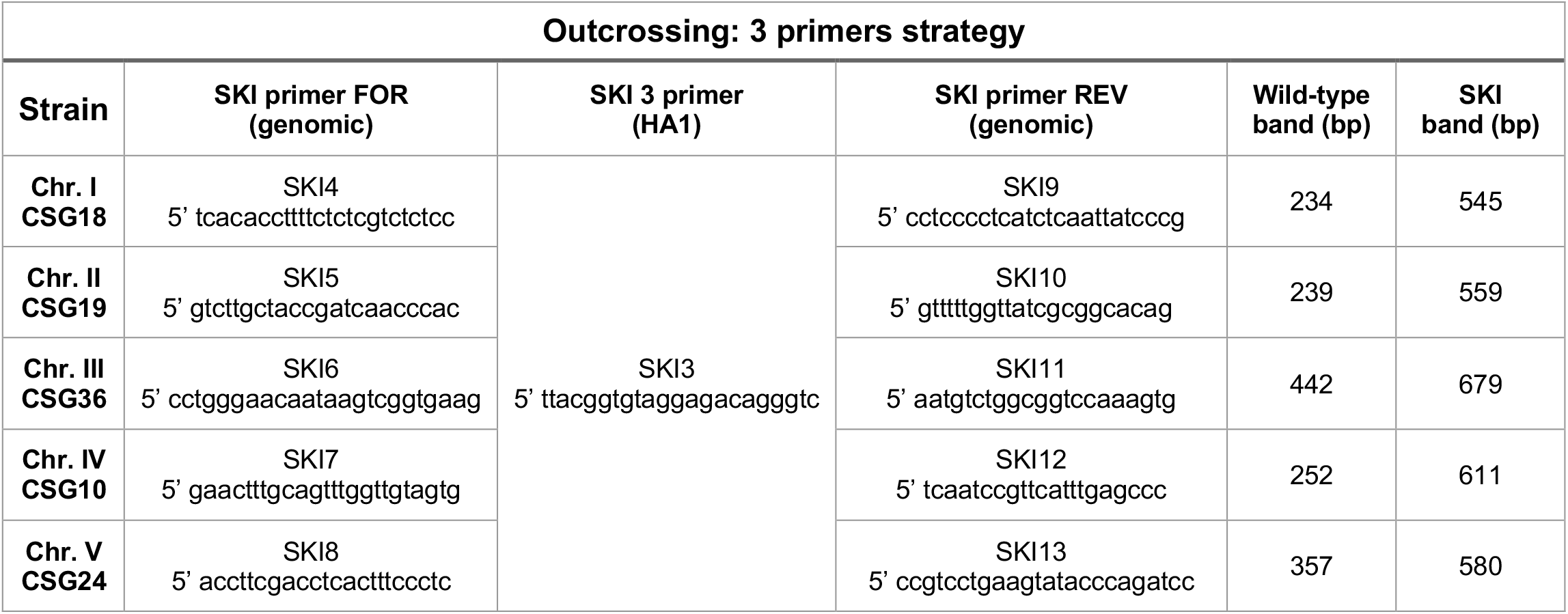

### Step 7 (optional)

The major predicted off-target site of the *dpy-10* crRNA is in an exon of R12E2.15. After outcrossing, to check potential off-target events from the *dpy-10* crRNA, we recommend using primers FOR (5’ gttgggtatgctcctccttgtg) and REV (5’ agaagactacatacgacggctgg). These primers amplify a fragment of 873 bp from the R12E2.15 gene, flanking the potential off-target sequence. Use this PCR product for sequencing with the same primers.

